# RNA-Protein Interaction Classification via Sequence Embeddings

**DOI:** 10.1101/2024.11.08.622607

**Authors:** Dominika Matus, Frederic Runge, Jörg K.H. Franke, Lars Gerne, Michael Uhl, Frank Hutter, Rolf Backofen

## Abstract

RNA-protein interactions (RPI) are ubiquitous in cellular organisms and essential for gene regulation. In particular, protein interactions with non-coding RNAs (ncRNAs) play a critical role in these processes. Experimental analysis of RPIs is time-consuming and expensive, and existing computational methods rely on small and limited datasets. This work introduces *RNAInterAct*, a comprehensive RPI dataset, alongside *RPIembeddor*, a novel transformer-based model designed for classifying ncRNA-protein interactions. By leveraging two foundation models for sequence embedding, we incorporate essential structural and functional insights into our task. We demonstrate RPIembeddor’s strong performance and generalization capability compared to state-of-the-art methods across different datasets and analyze the impact of the proposed embedding strategy on the performance in an ablation study.

## 1 Introduction

The discovery that 85% of the human genome is transcribed into ribonucleic acid (RNA), while only about 2% of these RNAs code for proteins (Birney et al., 2007; Consortium et al., 2012), has shifted our view of RNA from a mere translator between DNA and proteins to one of the most crucial cellular regulators. Although the functions of many non-coding RNAs (ncRNAs) remain unknown, it is widely acknowledged that their interactions with proteins are one of the driving forces for cellular functions, particularly in gene regulation and epigenetics (Oksuz et al., 2023; Statello et al., 2021; Mangiavacchi et al., 2023). However, experimental analysis of these interactions, e.g., via systematic evolution of ligands by exponential enrichment (SELEX) (Tuerk & Gold, 1990), is time-consuming and expensive. *In silico* methods capable of distinguishing between interacting and non-interacting RNA-protein pairs could significantly reduce these costs.

Deep learning-based methods recently broke new ground across a variety of applications in molecular research (Alipanahi et al., 2015; Ronneberger et al., 2015; Baek et al., 2021; Jumper et al., 2021; Baek et al., 2023). Additionally, meta-learning across tasks has demonstrated its potential to significantly improve the performance of deep learning models (Singh et al., 2019), particularly when labeled data is scarce - a common challenge in RNA-protein interaction (RPI) prediction tasks. While machine learning approaches to classify RPIs exist (Muppirala et al., 2011; Pan et al., 2016; Jain et al., 2018), the application of meta-learning strategies to leverage the diverse characteristics of various RNAs and proteins for this task remains largely unexplored. Furthermore, interactions typically hinge on structural features in addition to sequence information, which, for RNAs are not widely available at a large scale. An algorithm capable of accurately classifying ncRNA-protein interactions solely from sequence inputs and applicable across a wide range of interaction types would be highly valuable.

In this study, we introduce *RPIembeddor*, a novel and comprehensive approach for classifying ncRNA-protein interactions that addresses these challenges. Furthermore, we compile an extensive dataset of positive RPI entries from the RNAInter database (Kang et al., 2022) and enrich it with carefully generated negative examples, leveraging both sequence and structural features of the RNA and protein interactors. We employ two foundation models, RNA-FM (Chen et al., 2022) and ESM-2 (Lin et al., 2023), to generate embeddings for the RNA and protein sequences that are then used as inputs for the RPIembeddor.

Our main contributions are as follows:

- We build *RNAInterAct*^1^, an extensive dataset for ncRNA-protein interaction prediction, derived from the RNAInter (Kang et al., 2022) database. It comprises 73, 362 negative and 35, 852 positive interactions across 976 unique RNA families. To ensure rigorous evaluation, we meticulously curate two subsets, *TRinter* for training and *TSfam* for testing, with consideration to prevent any overlap in RNA families between the test and the training sets, thereby eliminating homology bias.
- We introduce *RPIembeddor*, a novel algorithm for ncRNA-protein interaction prediction that utilizes embeddings from two foundational models within an attention-based framework. Furthermore, we demonstrate its superior generalization capabilities when bench-marked against state-of-the-art models across various test sets, marking a significant advancement in the field.
- Through a comprehensive ablation study, we validate the usefulness of the selected embeddings, illustrating how they critically enhance model performance.

## 2 Related work

Due to the limited amount of experimentally derived RNA-protein complex structure data, machine learning methods typically rely on sequence information to predict RPIs (Bheemireddy et al., 2022). As an RNA-binding protein (RBP) can bind to many different RNAs with varying affinity depending on the presence and arrangement of specific structure and/or sequence recognition motifs in RNA, experimental interaction datasets for a specific protein can contain from thousands to tens of thousands RNA targets. These datasets are common and prevalently obtained from CLIP-seq experiments (Hafner et al., 2021), so most of the available computational methods use them to train protein-specific models to predict protein-binding sites on the given RNA sequences (Pan et al., 2019; Uhl et al., 2021). Consequently, these models depend on the availability of a sufficiently large interaction dataset for a protein of interest. However, out of the estimated 2,000 human RBPs (Brannan et al., 2016; Hentze et al., 2018; Liu et al., 2019), such datasets exist only for a few hundred, showcasing the need for alternative approaches. To study this vast space of unexplored RPIs, a particularly useful extension is approaches that predict whether any given RNA and protein interact based on their sequences. To date, only a limited number of methods have been developed for that particular task. To the best of our knowledge, these include RPIseq (Muppirala et al., 2011), IncPro (Lu et al., 2013), IPMiner (Pan et al., 2016), and XRPI (Jain et al., 2018). In the following, we will focus on XRPI and IPMiner, since they have been shown to outperform the previous two methods. A detailed description of the respective tools can be found in Appendix A.

## 3 Data

The foundation of our study on predicting RPIs is an extensive, meticulously compiled, and processed dataset. We provide a concise overview of our data pipeline in this section, with a more detailed description available in the Appendix B.

### Data Preparation

We use the RNA Interactome Database (RNAInter) (Kang et al., 2022) with over 47 million RNA interactions across 156 different species as the basis for our dataset. Among these, RPIs are particularly prominent, with 37,067,587 entries. Although RNAInter does not directly provide sequence information, we obtain it by cross-referencing different large-scale databases such as NCBI (Benson et al., 2012), UniProt (Consortium, 2019), and Ensembl (Cunningham et al., 2022). To enable informed negative interaction generation, we perform an extensive annotation using the Rfam (Kalvari et al., 2021) and Pfam (Mistry et al., 2020) databases, assigning RNA families and protein clans to each unique interactor. Our refinement process includes setting a length cutoff at 1024 nucleotides and amino acids, limiting the number of interactions per interactor to 150, and excluding mRNAs from the dataset. After careful curation of the negative examples by leveraging RNA family, protein clan and interactor’s category to ensure biological relevance, the *RNAInterAct* dataset encompasses a total of 122,217 interactions between ncRNAs and proteins with a 1:2 ratio of positive to negative interactions.

### Train and Test Sets

RNAInterAct serves as a foundation for the derived training and test sets, *TRinter* and *TSfam*, respectively. We strategically divide RNAInterAct based on the RNA families involved, ensuring no RNA family overlap between the sets. This approach is grounded in the understanding that RNA families consist of RNAs with conserved nucleotide sequences that share common structural features and typically, similar functions. Evaluating on such a test set ensures the assessment of the model’s true generalization capabilities, in stark contrast to the limited insights gained from random splits of training and test data.

In addition to evaluating *RPIembeddor* on the TSfam, we also assess it using the widely recognized RPI2825 dataset (Muppirala et al., 2011; Jain et al., 2018). This step allows us to examine the model’s ability to generalize across a distinct distribution of examples, as RPI2825 comprises exclusively positive interactions. Consistent with our methodology, we apply the same sequence length restriction of 1024 nucleotides/amino acids to this dataset. A comparative overview of the datasets utilized in our evaluations is presented in Table 1.

**Table 1:** Overview of the datasets used in this work.

| Feature | TRinter | TSfam | RPI2825 |
| --- | --- | --- | --- |
| Unique RNA Families | 976 | 172 | N/A |
| Unique Protein Clans | 152 | 152 | N/A |
| Positive Interactions | 35,852 | 4,887 | 871 |
| Negative Interactions | 73,362 | 8,116 | 0 |
| Total Interactions | 109,214 | 13,003 | 871 |

## 4 Method

In this section, we detail *RPIembeddor*, our proposed approach for RPI classification.

### Embeddings

The centerpiece of our model involves leveraging RNA and protein embeddings from pre-trained models to incorporate structural and functional insights into our interaction prediction task. For RNA sequences, we employ the transformer-based RNA Foundation Model (RNA-FM) (Chen et al., 2022), which was trained on a massive corpus of 23 million unlabeled non-coding RNAs from RNAcentral (Consortium et al., 2017) using self-supervised learning. For protein sequences, we utilize Evolutionary Scale Modeling 2 (ESM-2) (Lin et al., 2023), a transformer-based language model trained on the UniRef database (Suzek et al., 2014) that specializes in predicting protein folding from amino acid sequences. Contrasting with the well-known AlphaFold (Jumper et al., 2021), ESM-2 operates without the need for multiple sequence alignments (MSAs), making it a more fitting option for our requirements. We opt for the 30-layer version of ESM-2 with 150 million parameters to match RNA-FM’s embedding size of 640. Given that RNA-protein interactions hinge critically on the structures and functional characteristics of the molecules involved, the combined use of embeddings from these two models offers a comprehensive view of potential interaction sites and mechanisms, promising performance improvement on the RPI classification task.

### Model

After processing the inputs with RNA-FM and ESM-2, we obtain two embeddings, each of size *N ×*640 with *N* being the input sequence length. These embeddings serve as an input for the RPIembeddor. To address the possibility of varying lengths of RNA and protein sequences while ensuring compatibility, we implement two parallel feed-forward layers that normalize the size of the input embeddings. Subsequently, the embeddings undergo processing in an encoder layer designed to treat them symmetrically, ensuring they have equal influence on the model’s final output probability. This symmetrical processing is crucial as it allows our model to dynamically focus on specific parts of the sequences that are most relevant for predicting interactions, leveraging the strengths of the attention mechanism. By doing so, attention facilitates the model’s ability to capture complex dependencies between RNA and protein sequences. The resulting latent representations are concatenated and processed through a series of feed-forward layers, culminating in a linear layer with a sigmoid activation function to produce output class probabilities. The architectural choices result in the model size of 1.4M parameters. For the task of RPI classification, we employ a binary loss function and optimize the model using a combination of linear warm-up and cosine annealing strategies with the AdamW optimizer (Loshchilov & Hutter, 2019). For a detailed overview of RPIembeddor’s architecture and hyperparameters, please refer to Appendix C.1.

## 5 Experiments

To evaluate *RPIembeddor*’s efficacy, we conduct two sets of experiments. First, we compare its performance against state-of-the-art methods, namely IPMiner (Pan et al., 2016) and XRPI (Jain et al., 2018), on both the *TSfam* and the RPI2825 datasets. Secondly, through an ablation study, we analyze the impact of embeddings on model performance. For both experiments, we report performance metrics such as binary precision, recall, F1-score, and accuracy, and for the first set additionally include plotting Receiver Operating Characteristic (ROC) curves.

### 5.1. Benchmarking on TSfam and RPI2825

#### Setup

For a robust evaluation, RPIembeddor is trained on TRinter using three distinct random seeds. Its results are aggregated to report the average performance alongside the standard deviation. We do not retrain the competing models, as the authors do not provide comprehensive instructions to do so, and we use their publicly accessible versions.^2^ ^3^ All models are evaluated on two distinctive datasets, as detailed in Table 1. We present the results in Table 2.

**Table 2:** Mean performance and standard deviation across three seeds of the RPIembeddor in comparison to state-of-the-art models.

| Model | TSfam |  |  |  | RPI2825 |  |  |  |
| --- | --- | --- | --- | --- | --- | --- | --- | --- |
|  | Prec. | Rec. | F1 | Acc. | Prec. | Rec. | F1 | Acc. |
| RPIembeddor | **0.550**<br>**$\pm 0.010$** | 0.627<br>$\pm 0.017$ | **0.586**<br>**$\pm 0.013$** | **0.667**<br>**$\pm 0.009$** | **1.0**<br>**$\pm 0.0$** | 0.667<br>$\pm 0.085$ | 0.8<br>$\pm 0.049$ | 0.667<br>$\pm 0.085$ |
| IPMiner | 0.357 | 0.375 | 0.366 | 0.512 | **1.0** | 0.107 | 0.193 | 0.107 |
| XRPI | 0.375 | **0.909** | 0.531 | 0.398 | **1.0** | **0.982** | **0.991** | **0.982** |

#### TSfam

We observe that RPIembeddor demonstrably outperforms competing models, achieving an F1-score of 0.586 (±0.010) and an accuracy of 0.667 (±0.009). Specifically, our model correctly classifies 2,971 out of 4,887 positive interactions and 5,586 out of 8,116 negative ones. In comparison, IPMiner predicts 1,830 true positives and 4,826 true negatives. Notably, XRPI exhibits a significant bias towards positive classifications, predicting approximately 91% of interactions (11,832 out of 13,003) as positive, despite the dataset comprising roughly 62% negative examples. We illustrate Receiver Operating Characteristic (ROC) curves for all three models in Figure 1.

**Figure 1.**
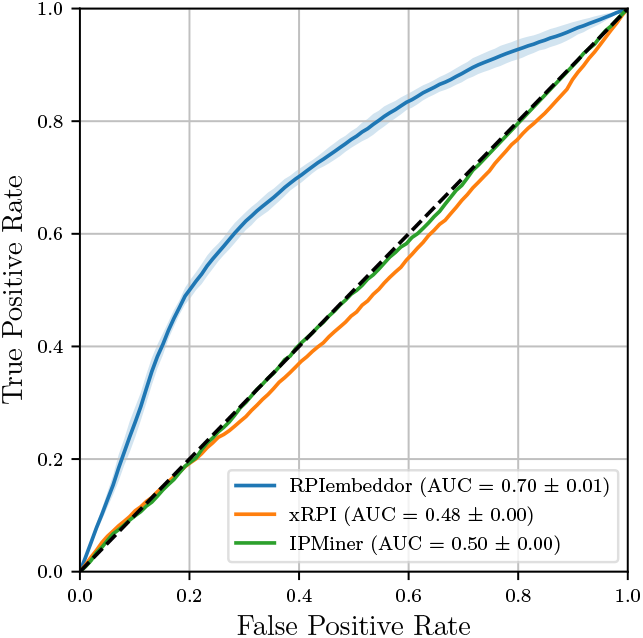
Receiver Operating Characteristics comparison on TSfam.

**Figure 2.**
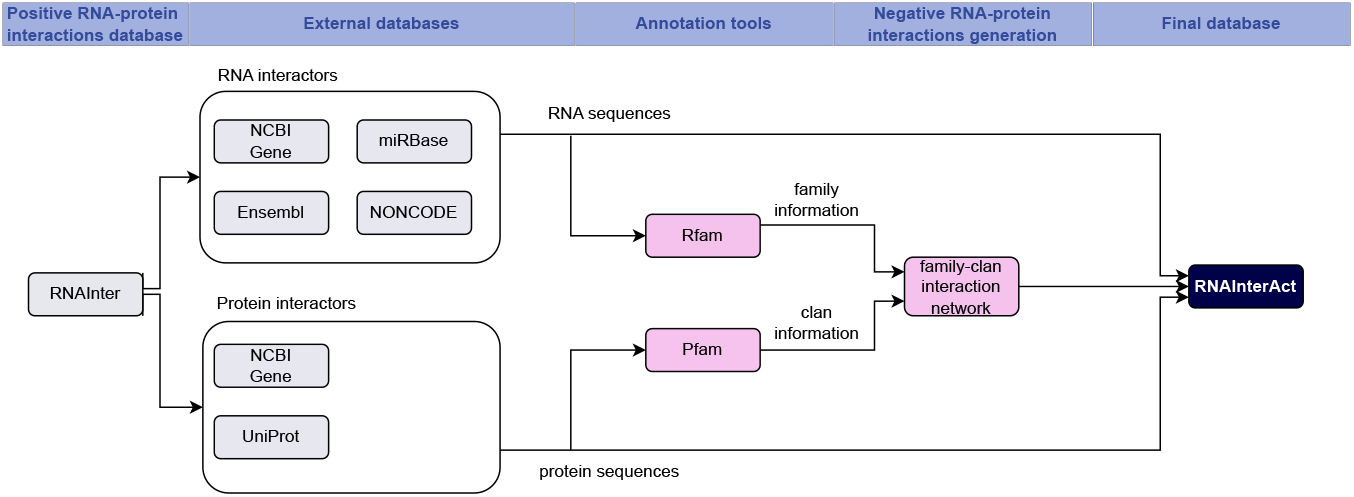
Data curation and processing pipeline.

**Figure 3.**
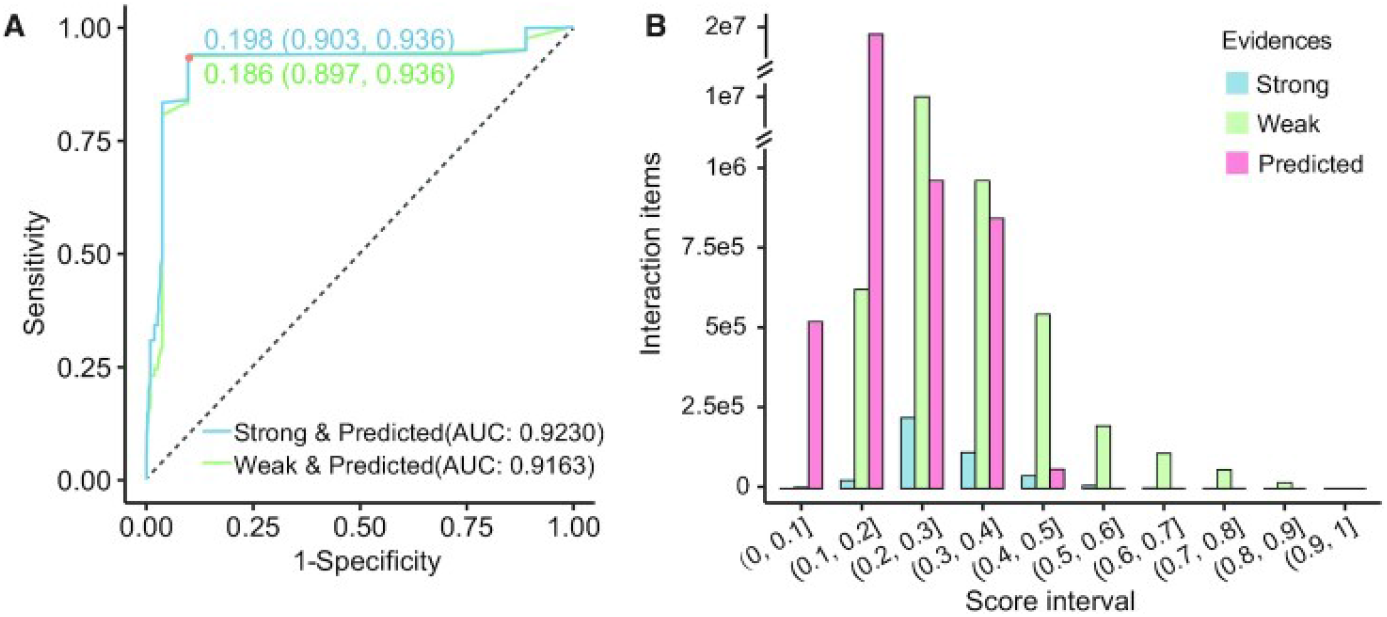
Evaluation of confidence scores. (A) ROC for distinguishing between experimental and predicted interactions. (B) Interaction number for each score interval. Plots are adapted from (Kang et al., 2022).

**Figure 4.**
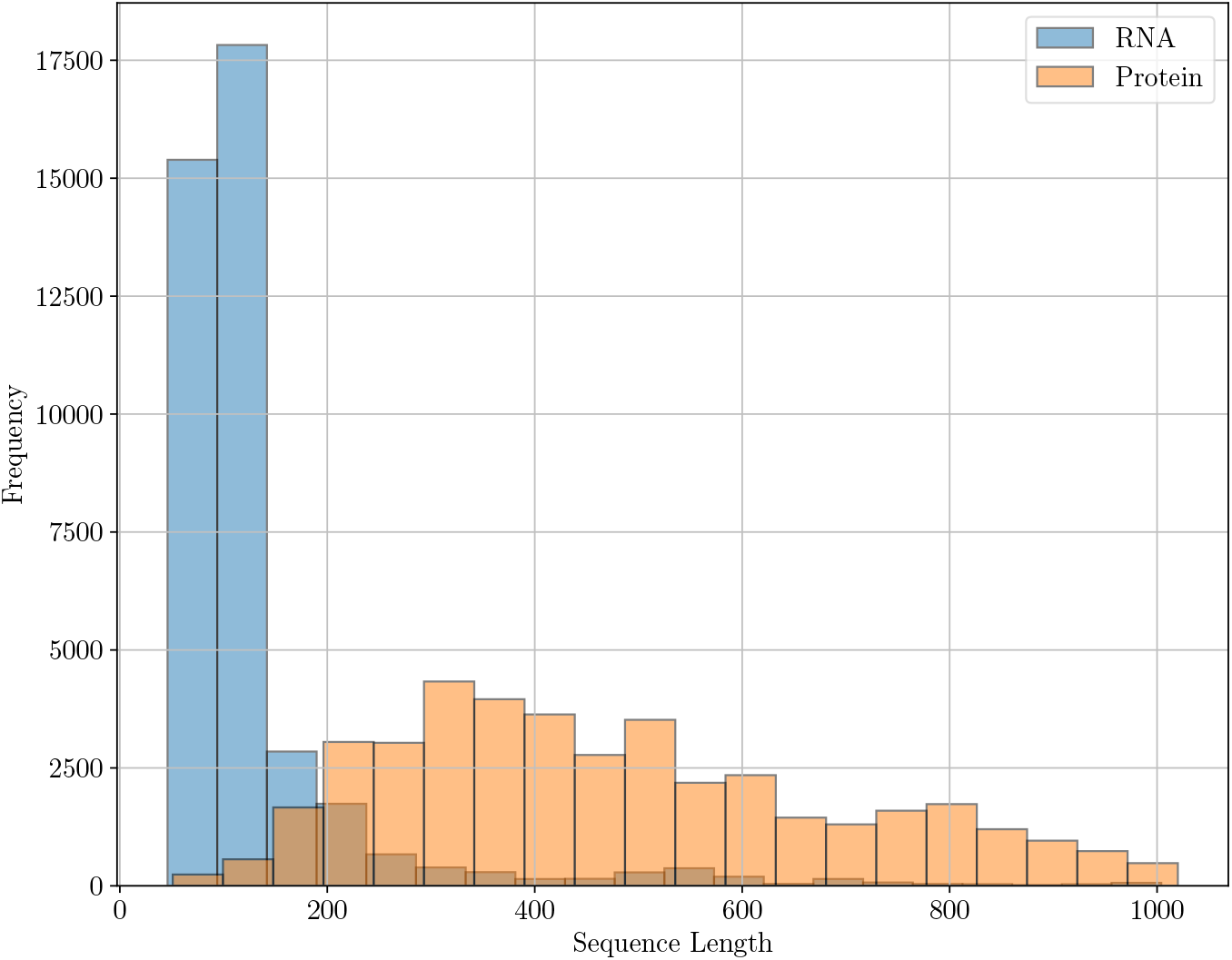
Sequence length distribution.

**Figure 5.**
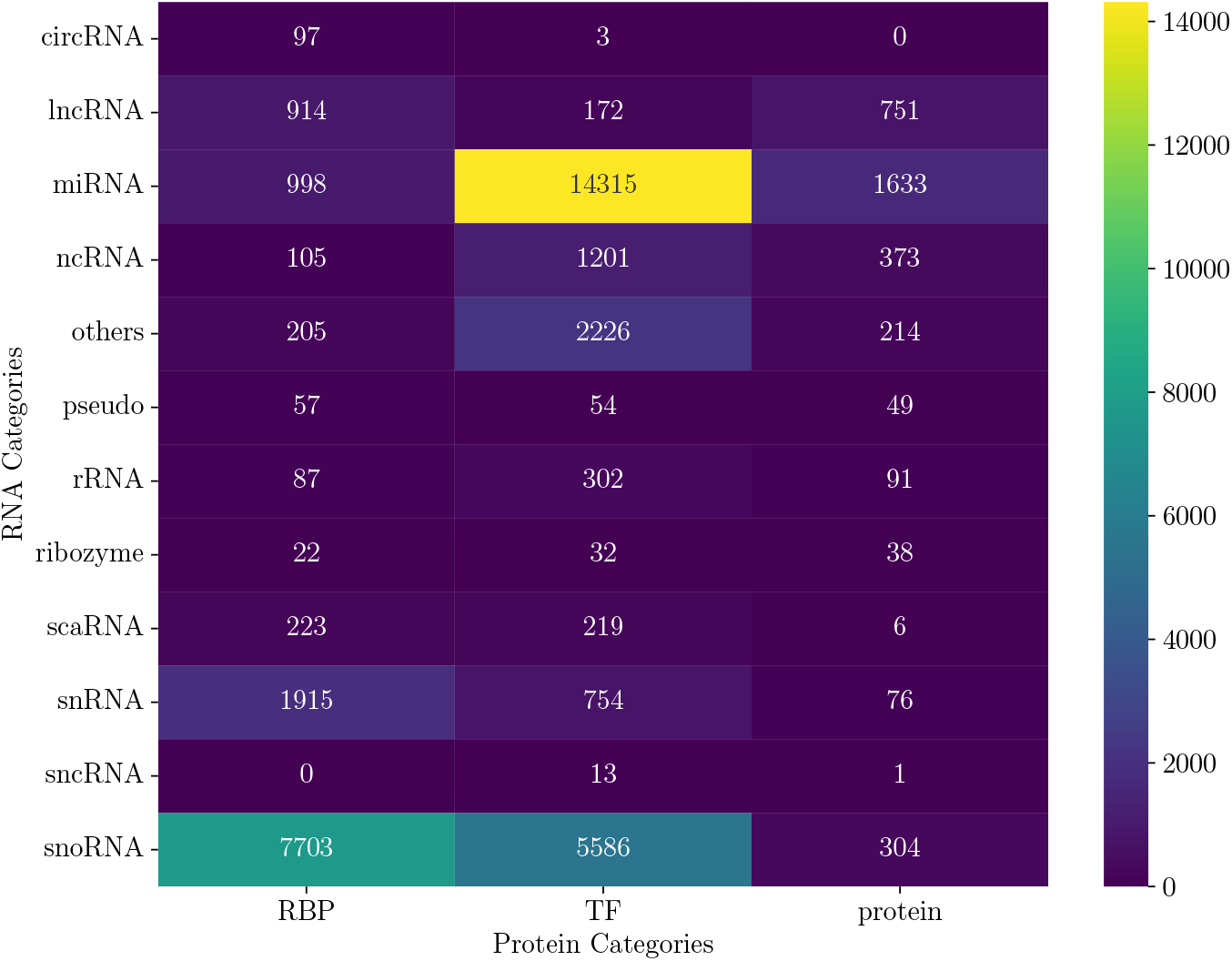
Positive interactions per RNA/protein category heatmap.

**Figure 6.**
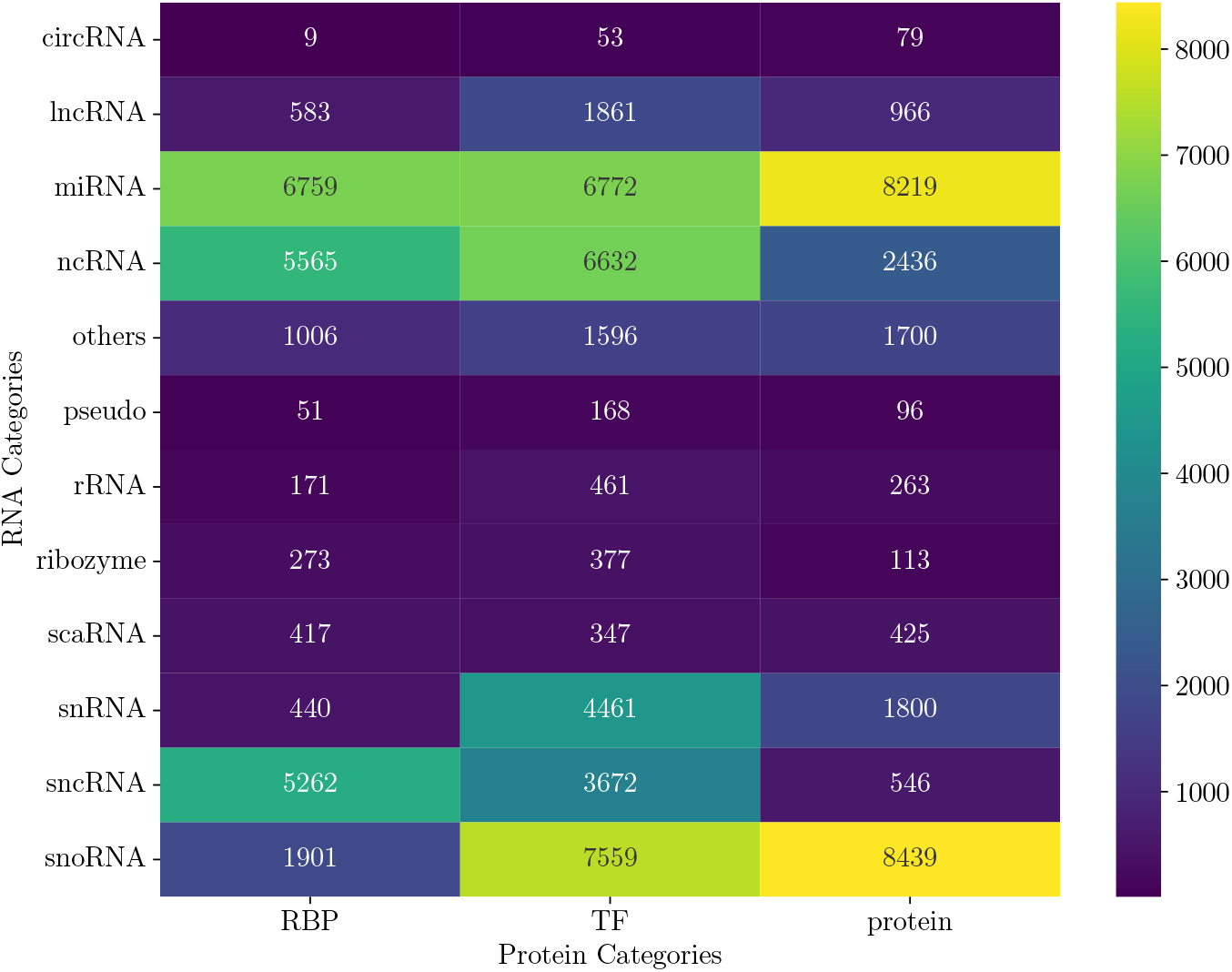
Negative interactions per RNA/protein category heatmap.

**Figure 7.**
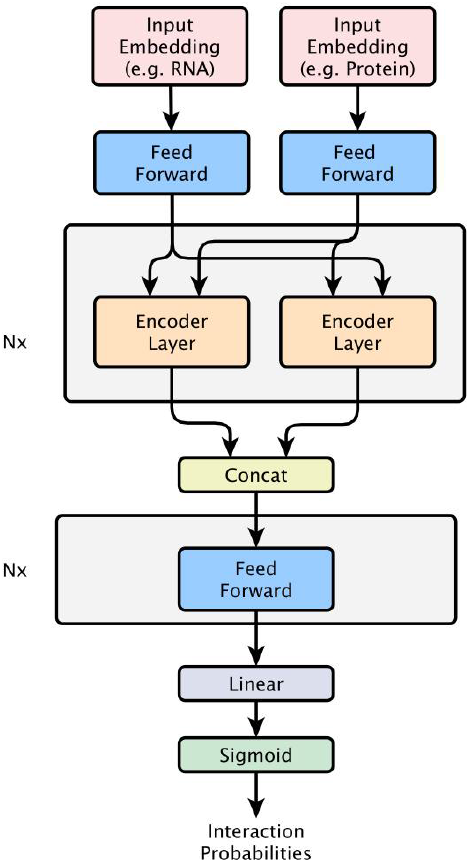
RPIembeddor architecture overview.

**Figure 8.**
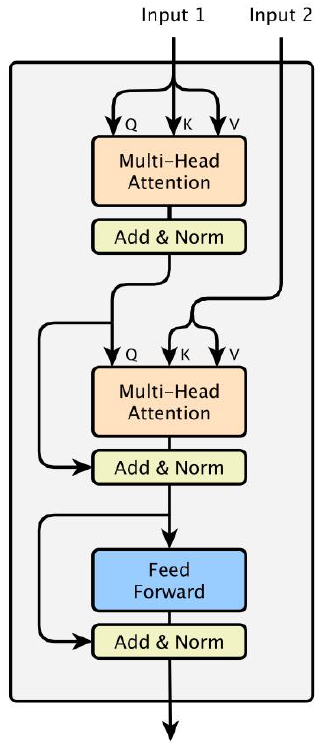
Encoder layer.

#### RPI2825

The RPI2825 dataset comprised exclusively of positive interactions presents a challenge for RPIembeddor. Despite this, our model demonstrates robust generalization, achieving the second-best F1-score of 0.8 (±0.049), giving in to XRPI with an impressive F1-score of 0.991. However, it is important to note that XRPI was trained on the RPI2825 dataset—using the original 2825 entries without the modifications described in Section 3. Our analysis suggests that RPIembeddor’s performance is not merely a reflection of the training data distribution, as it effectively generalizes to unseen data distributions such as positive interactions-only RPI2825. This underscores the robustness and versatility of our model.

### 5.2 Ablation Study

### Setup

We conduct the analysis of embeddings’ impact on model performance in two parts: (i) by replacing either the protein or RNA input embedding with a random embedding of the same size to investigate if both embeddings contribute equally to performance, and (ii) by replacing both input embeddings with one-hot encodings of the input sequences to assess whether our embeddings are superior to simpler representations in capturing the relationships between the RNA and protein sequences. We train one distinct model for each configuration under RPIembeddor’s architecture: *random-protein* utilizing only ESM-2 protein embedding; *RNA-random* using only the RNA-FM RNA embedding; and *one-hot*, where both RNA and protein sequences are represented using one-hot encodings. Each model is trained on TRinter and evaluated on TSfam using three unique random seeds to ensure the reliability of our results. We report binary precision, recall, F1-score, and accuracy for these models in Table 3 and compare them to the original RPIembeddor’s performance, which utilizes both ESM-2 and RNA-FM embeddings.

**Table 3:** Results of the ablation study.

| Model | Prec. | Rec. | F1 | Acc. |
| --- | --- | --- | --- | --- |
| RPIembeddor | **$0.563 \pm 0.019$** | **$0.659 \pm 0.071$** | **$0.605 \pm 0.019$** | **$0.678 \pm 0.009$** |
| One-Hot | $0.0 \pm 0.0$ | $0.0 \pm 0.0$ | $0.0 \pm 0.0$ | $0.624 \pm 0.0$ |
| Random-Protein | $0.0 \pm 0.0$ | $0.0 \pm 0.0$ | $0.0 \pm 0.0$ | $0.624 \pm 0.0$ |
| RNA-Random | $0.0 \pm 0.0$ | $0.0 \pm 0.0$ | $0.0 \pm 0.0$ | $0.624 \pm 0.0$ |

**Table 4:**
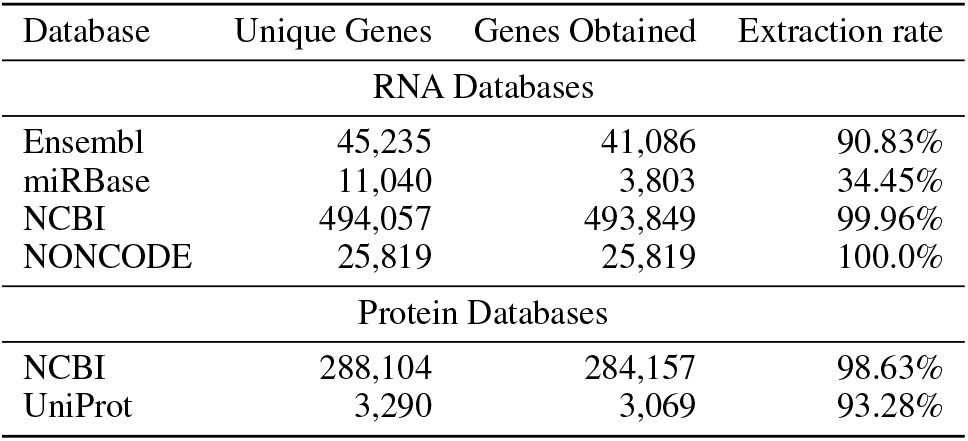
RNAInter external databases statistics.

| Database | Unique Genes | Genes Obtained | Extraction rate |
| --- | --- | --- | --- |
| RNA Databases |  |  |  |
| Ensembl | 45,235 | 41,086 | 90.83% |
| miRBase | 11,040 | 3,803 | 34.45% |
| NCBI | 494,057 | 493,849 | 99.96% |
| NONCODE | 25,819 | 25,819 | 100.0% |
| Protein Databases |  |  |  |
| NCBI | 288,104 | 284,157 | 98.63% |
| UniProt | 3,290 | 3,069 | 93.28% |

**Table 5:** Summary of family and clan annotation.

| Type | Unannotated | Annotated | Families/Clans |
| --- | --- | --- | --- |
| RNA | 69,043 | 7,847 | 1,148 |
| Protein | 38,026 | 26,575 | 152 |

**Table 6:**
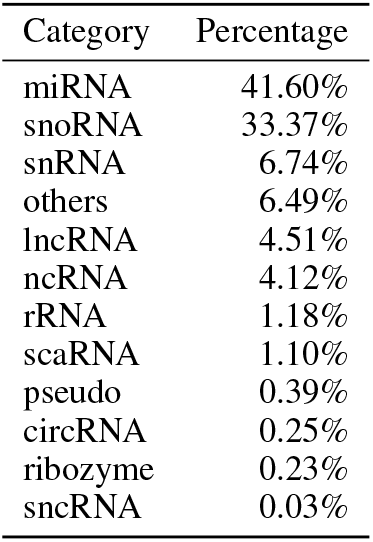
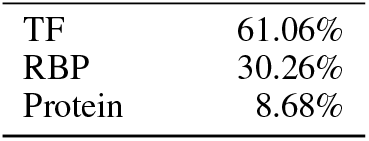
Breakdown of RNA (left) and protein (right) categories comprising the dataset.

| Category | Percentage |  | Category | Percentage |
| --- | --- | --- | --- | --- |
| miRNA | 41.60% |  | TF | 61.06% |
| snoRNA | 33.37% |  | RBP | 30.26% |
| snRNA | 6.74% |  | Protein | 8.68% |
| others | 6.49% |  |  |  |
| lncRNA | 4.51% |  |  |  |
| ncRNA | 4.12% |  |  |  |
| rRNA | 1.18% |  |  |  |
| scaRNA | 1.10% |  |  |  |
| pseudo | 0.39% |  |  |  |
| circRNA | 0.25% |  |  |  |
| ribozyme | 0.23% |  |  |  |
| sncRNA | 0.03% |  |  |  |

**Table 7:** Overview of unique sequences, families, and clans of the dataset.

| Proteins | RNA | Pfam clans | RNA families |
| --- | --- | --- | --- |
| 1,308 | 4,169 | 152 | 1,148 |

### Results

RPIembeddor in all three configurations consistently behaves as a negative classifier, predicting only negative examples. This results in binary precision, recall, and F1-score of 0.0 as no positive examples as classified correctly (true positives are 0). An accuracy of 0.624 merely reflects the data distribution in TSfam - 7,991 out of 12,807 samples are negatives, correctly identified as such. This underscores the significance of ESM-2 and RNA-FM embeddings as meaningful representations of protein and RNA sequences, respectively. These embeddings carry structural and functional information critical for correct classification. Furthermore, the results affirm that both embeddings are essential for the model to perform effectively, aligning with the intuition that RPIs depend crucially on both the protein and RNA involved.

### 6 Conclusion

Our work introduces *RPIembeddor*, a transformative approach to RNA-protein interaction (RPI) prediction that harnesses the power of embeddings from two foundational models. Using a meta-learning strategy to learn RPIs across different RNA and protein types, our method outperforms existing methods while generalizing to unseen data distributions. We believe that our approach bears great potential for future RPI prediction endeavors and we support this research by making our new dataset *RNAInterAct* publicly available. Acknowledging the limitations tied to foundational model dependencies and sequence length constraints, our future directions include exploring alternative embeddings, e.g., from RNA structure models like the RNAformer (Franke et al., 2023) or other foundation models, to extend the applicability of our approach.

## 7 Acknowledgments

This research was funded by the German Research Foundation (DFG) under SFB 1597 (SmallData), grant no. 499552394, and through grant no. 417962828. The authors acknowledge support by the state of Baden-Württemberg through bwHPC and the German Research Foundation (DFG) through grant INST 35/1597-1 FUGG and INST 39/963-1 FUGG. Dominika Matus further acknowledges funding by the Konrad Zuse School of Excellence in Learning and Intelligent Systems (ELIZA) through Master’s in AI scholarship. Finally, we acknowledge funding by the European Union via ERC Consolidator Grant DeepLearning 2.0, grant no. 101045765. Views and opinions expressed are however those of the author(s) only and do not necessarily reflect those of the European Union or the European Research Council. Neither the European Union nor the granting authority can be held responsible for them.

## A Related Work

In this section, we describe existing computational methods of the RNA-protein interaction (RPI) prediction based on sequence information.

### RPIseq

RPIseq (Muppirala et al., 2011) predicts RPIs by extracting sequence-based features and using these as inputs to two machine learning models: Random Forest (RF) and Support Vector Machine (SVM). The primary dataset, RPI2241, is derived from the Protein-RNA Interface Database (PRIDB) and contains only positive samples. To balance the dataset for training, negative samples are generated through random pairing, guided by sequence identity. It provides an online interface for running inference on either of the models ^4^.

### IncPro

IncPro (Lu et al., 2013) calculates interaction score between long non-coding RNA (lncRNA) and protein based on the features derived from the physicochemical properties of amino acids and nucleotides. It generates a scoring matrix reflecting the propensity of each protein residue to interact with each RNA residue and then uses a statistical model to calculate the overall likelihood of the pair interacting. The tool is no longer available under the address reported by the authors.

### IPMiner

IPMiner (Pan et al., 2016) uses stacked autoencoders that extract features from the non-coding RNA (ncRNA) and protein sequences that are then forwarded to an RF classifier, with stacked ensebling employed to integrate multiple model outputs. It uses experimentally derived interaction pairs from Protein Data Bank (PDB) and NPInter2.0 and generates the negative samples through random pairing. Model can be accessed through the github repository ^5^.

### XRPI

XRPI (Jain et al., 2018) utilizes the Extreme Gradient Boosting (XGBoost) algorithm to predict RPIs. High-resolution structural data is derived from PDB in order to analyze amino acid interaction propensities and determine the optimal length of sequence windows in RNAs, the features later applied to the input sequences data during the training phase. Similarly to RPIseq, it uses RPI2825 dataset for training and evaluation. The negative interactions are generated by “random jumbling” of RNA and proteins under sequence similarity condition. Model is accessible via authors department website ^6^.

## B Data

The complete data processing pipeline is presented in Figure 2.

### B.1 The RNA Interactome Database

The RNA Interactome Database (RNAInter) (Kang et al., 2022) is a specialized resource in the field of molecular biology that houses an extensive collection of over 47 million RNA interactions of various types coming from 156 different species. Among these, RNA-protein interactions (RPIs) are particularly prominent, with slightly over 37 million entries. This significant volume of data underlines the importance of the complex interplay between RNA and proteins within biological systems.

RNAInter v4.0, as utilized in this project, expands on its predecessor, RNAInter v3.0, through extensive literature mining and the integration of external databases with interactions sourced from experimental evidence or computational prediction. Each entry within RNAInter v4.0 is assigned a confidence score ranging from 0 to 1, reflecting a comprehensive evaluation based on three key factors: the reliability of experimental evidence, community trust, and the specificity of cells or tissues involved. The score distribution for all present interactions as presented in Figure 3. It is visibly right-skewed, with the majority of entries assigned either “weak” or “predicted” evidence categories. However, despite this skewness indicating a potential abundance of lower-confidence interactions, the database’s inclusivity in capturing a wide range of interactions, including less-studied ones, presents valuable opportunities for exploratory research.

Among the databases contributing to the RPI data are LncTarD (Zhao et al., 2022), with experimentally validated interactions; oRNAment (Bouvrette et al., 2020), a repository for computationally predicted interactions; and NPInter v4.0 (Teng et al., 2020), which includes both types. Together, these sources provide a comprehensive dataset of 37,067,587 RPI interactions, enriched with details such as species, target regions, tissues or cell lines, and homology interactions. While the entries do not explicitly include sequence information, each interactor is linked to an external database, facilitating the retrieval of such critical data for our project.

### B.2 Sequence data

To complete the interaction data from the RNA Interactome Database (RNAInter) with essential sequence information, we access databases linked to each interactor in a single entry. Protein sequences are sourced from NCBI (Benson et al., 2012), and UniProt (Consortium, 2019), while for RNA sequences, our sources include miRBase (Kozomara et al., 2019),Ensembl (Cunningham et al., 2022), NONCODE (Fang et al., 2018), and NCBI. Besides the sequence, we also query for sequence length and specific IDs. These IDs are crucial for linking each sequence to its corresponding entry in the RNAInter database. For a detailed overview of our data extraction rates relative to RNAInter contents and the total number of sequences compiled, please refer to Table 4.

During our data processing, we exclude any entry that lacks either sequence information or necessary IDs. We also eliminate duplicates based on the ID. This step is important because the same sequence might appear in multiple databases and be associated with different interactions in the RNAInter database. Finally, owing to the input length limitations of the foundation models employed for embedding generation, we set a maximum sequence length of 1024 for both RNA and protein molecules. That results in a total number of 38,026 protein entries and 69,043 RNA entries.

For the subsequent stages of annotation and embedding generation, our focus shifts to the individual protein and RNA sequences, rather than the interactions they form.

### B.3 Annotations

Our approach to generating negative interactions, as detailed in Section B.5, builds upon the notion of similarity between different RNA or protein sequences. An established way of expressing that is through family annotation as it is done in Pfam and Rfam databases, for protein and RNA sequences respectively.

The Pfam database (Mistry et al., 2020) represents a comprehensive collection of protein families, each characterized through multiple sequence alignments (MSA) and hidden Markov models (HMMs) that help identify function regions within proteins, called domains, and group them based on shared characteristics. Additionally, Pfam introduces a higher level of classification known as ‘clans’, which groups together families that share a single evolutionary origin, as confirmed by similarities in sequence, structure, and profile-HMMs. For our project, we utilized Pfam 36.0, which comprises 20,795 entries and 659 clans.

The Rfam database(Kalvari et al., 2021) comprises a curated collection of non-coding RNA (ncRNA) families. Each family in this database is characterized by a multiple sequence alignment and a consensus secondary structure, accompanied by a covariance model that aids in the annotation of new family members. This classification, which includes structural information, is particularly crucial for ncRNAs. Unlike protein-coding genes, ncRNAs often exhibit significant functional similarities linked to their secondary structures, even when their primary sequences show little resemblance. Such considerations are vital for an approach that utilizes primary sequence data. In our project, we access Rfam 14, encompassing 4,170 families.

We scan each unique protein sequence against the Pfam database and for recognized entries, assign a clan name. Similarly, each unique RNA sequence is scanned against the Rfam database using Infernal (Nawrocki & Eddy, 2013) tool, and, if found, annotated a family name. Results concerning the number of sequences are stored in Table 5.

### B.4 Positive interactions dataset

After gathering all necessary sequence information for generating negative interactions, we then focus on creating a dataset of positive interactions. We cross-reference the annotated RNA and protein sequences, as detailed in Table 5, with the RNA Interactome Database (RNAInter). Our criterion for inclusion is an overlap of sequence IDs between our dataset and RNAInter. This process yields 488,184 interactions, which represents approximately 1% of the original RNA-protein interactions (RPI) data entries in RNAInter. Such a reduction is anticipated, given several limiting factors we applied: sequence length restrictions, the requirement for family or clan annotations, and variable extraction rates from the linked databases.

In the process of refining our dataset, we undertake several critical steps of analysis and filtering. Initially, we examine the category information of both RNA and protein sequences. From the RNA dataset, we exclude all mRNA sequences and categories that are unspecified. Despite the significant role of mRNAs in RPIs, our methodology is constrained by the RNA-FM foundation model used for generating RNA embeddings. This model is exclusively pre-trained on ncRNAs, and including mRNAs could potentially compromise the quality of the embeddings, thereby adversely affecting prediction performance.

While we acknowledge the existence of other foundation models like CodonBERT (Li et al., 2023), UTR-LM (Chu et al., 2023), and 3UTRBERT (Yang et al., 2023), which are specialized for coding sequences (CDS), 5’ untranslated regions (UTRs), and 3’UTR mRNAs respectively, their integration presents practical challenges. Given that some of the interactions in the RNAInter include genomic context, the deployment of additional foundation models would be computationally expensive and impractical for inference purposes, thus limiting the utility of our tool. Moreover, the Rfam database, as discussed in Section B.3, does not differentiate between various mRNA families, a distinction that is crucial for our approach for generating negative examples.

Consequently, after careful consideration, we remove 25,010 mRNA interactions and an additional 186 invalid RNA sequences from our dataset. The protein categories are limited to three types: transcription factors (TF), RNA-binding proteins (RBP), and general proteins. Given this concise categorization, we decide to retain all protein interactors. Following the removal of duplicates, our final dataset comprises 462,988 interactions.

In the final stage of data preparation, we impose a limit of 150 interactions per interactor. This threshold was determined through thorough analysis as an optimal balance, allowing us to maintain a reasonable number of interactions while preventing the over-representation of certain interactors in the dataset. As a result of this limitation, our collection of positive interactions is now 40,744. Given that we intend to generate two negative interactions per one positive, the dataset will eventually triple in size. This expansion ensures that even after the significant reduction in interactions, the dataset remains robust enough to support the effective performance of our model. We present the interactors’ contributions per category in Table 6.

The sequence length distributions, as shown in Figure 4, indicate that RNA sequences in the RNAInter database are typically much shorter than protein sequences, with the majority falling within the range of approximately 50 to 250 nucleotides (nts). This observation validates our sequence length restriction criteria, confirming that it did not excessively exclude sequences from our dataset. On the other hand, the protein sequence length distribution appears to be normally distributed with a slight right skew. This reflects the tendency of proteins to form longer amino acid (AA) chains, corroborating their nature as generally larger molecules.

Based on the data presented in Table 7, we observe that the RNA family might be a relatively weak indicator of similarity among different RNA sequences, as it appears that, on average, each family groups together about three sequences. In contrast, Pfam clans seem to offer a more distinct separation between protein sequence clusters, with an average of approximately nine protein sequences per clan. These annotations are crucial in our method for generating negative examples, where finding relations between interacting clans and families is key.

Finally, we examine the various types of interactions within our dataset, focusing on the categories of the interactors. MicroRNAs (miRNAs) dominate the RNA interactor category, which is biologically plausible given their crucial role in gene expression regulation; hence, the high number of miRNA-transcription factor (TF) pairs observed. Similarly, the small nucleolar RNA (snoRNA) category is prominently represented on the heatmap, suggesting numerous interactions related to small nucleolar ribonucleoproteins (snoRNPs), which consist of snoRNAs and associated proteins. Our dataset exhibits a wide range of other interaction types as shown in Figure 5.

### B.5 Negative interactions dataset

As discussed previously, the conventional method of generating negative interactions involves randomly selecting RNA and protein sequences not present in the positive samples. However, this approach could result in a dataset with high levels of noise, as it essentially relies on arbitrary selection without specific biological rationale.

In our proposed approach, we focus on making more informed decisions based on family/clan information and, where relevant, interactor categories. Initially, we established a network of family-clan interactions. For each RNA family member, we analyze the positive interactions dataset to identify its protein interactors, linking the RNA family to the clans of these proteins. Additionally, we annotate each RNA family with information about the protein categories that interact with its members, and each clan with data on the RNA categories interacting with its protein members. This methodology enables a broader scale analysis of RNA-protein interactions, moving beyond individual sequence analysis to understand the behavior of similar RNA and protein groups.

For each positive interaction, we generate two corresponding negative interactions, utilizing the family-clan interaction data and information on interacting categories. In the first negative interaction, we retain the original RNA interactor but pair it with a protein from a clan that does not interact with the RNA’s family and, if possible, is not part of the interacting categories. Similarly, for the second negative interaction, we keep the original protein interactor and pair it with an RNA from a family that does not interact with the protein’s clan and is not part of the interacting category. This approach ensures more accurate modeling of potential interactions, avoiding arbitrary pairings and focusing instead on biologically plausible non-interactions.

After merging the newly generated negative interactions dataset with the existing positive interactions dataset, we obtain a comprehensive dataset with 122,217 entries that is well-suited for binary classification tasks.

Figure 6 presents the distribution of the generated negative interactions per RNA/protein category.

## C Methods

### C.1 Model

In the development of RPIembeddor, we drew significant inspiration from transformer-based architectures, which have revolutionized the field of natural language processing (NLP) and, more recently, demonstrated their applicability and effectiveness within molecular biology (Jumper et al., 2021; Lin et al., 2023; Chen et al., 2022; Alipanahi et al., 2015; Brandes et al., 2022). The core principles of transformers(Vaswani et al., 2017), particularly their ability to capture long-range dependencies through self-attention mechanisms, are exceptionally well-suited to understanding the complex, sequence-based interactions characteristic of RNA and proteins.

To tailor RPIembeddor for the domain of RNA-protein interaction prediction, we carefully calibrated its architecture, hyperparameters and optimization strategy. The model’s backbone consists of an encoder with feature vectors dimensionality set to *d*_*model*_ = 256 and employs a multi-head attention mechanism with *n*_*head*_ = 2 to efficiently process sequence information. The incorporation of a single encoder layer, coupled with a deep feedforward network comprising 20 layers, strikes a balance between model complexity and interpretability.

For the optimization strategy we opted for cosine annealing scheduler with the AdamW optimizer(Loshchilov & Hutter, 2019), and introduced a warm-up phase for the first 1,000 training steps, initializing scheduler with a learning rate of 0.001. This warm-up phase helps avoid too fast convergence to suboptimal solutions by gradually increasing the learning rate. Following this, the cosine annealing scheduler reduces the learning rate over time, improving the model’s fine-tuning and generalization capabilities.

To prevent overfitting, we applied weight decay, setting it at 0.1, and used a dropout rate of 0.3.

This hyperparemeter configuration was applied consistently for 90 epochs during the training phase with a batch size of 64 across three different seeds.

### Footnotes

1 RNAInterAct is publicly available at https://ml.informatik.uni-freiburg.de/research-artifacts/RNAInterAct/RNAInterAct.tar.gz

2 XRPI: https://universe.bits-pilani.ac.in/goa/aduri/xRPI

3 IPMiner: https://github.com/xypan1232/IPMiner

4 http://pridb.gdcb.iastate.edu/RPISeq

5 https://github.com/xypan1232/IPMiner

6 https://universe.bits-pilani.ac.in/goa/aduri/xRPI

